# Endocytosis of PEGylated polymeric mesoscale nanoparticles is dynamin- and macropinocytosis-dependent

**DOI:** 10.64898/2026.01.22.701067

**Authors:** Adnan Arnaout, Poojaa Jayanthi Venugopal, Pratyusha Ghosh, Ryan Williams

## Abstract

Nanotechnology is rapidly transforming medicine by enabling versatile platforms for targeted delivery, controlled release, and intracellular transport of therapeutic payloads. Polymeric mesoscale nanoparticles (MNPs) are 300 to 500 nm in diameter with a PEGylated surface that exhibit unique renal tropism, specifically toward renal tubular epithelial cells. Despite their well-described therapeutic applications and route of localization to the tubules, we do not yet understand their physicochemical stability and cellular internalization mechanisms. In this study, we investigated the stability of MNPs under stress conditions by subjecting them to repeated freeze-thaw cycles and varying storage conditions to evaluate the effects on particle size and polydispersity index. MNPs demonstrated negligible changes in size and PDI up to 4 freeze-thaw cycles. We found that both empty and dye-loaded MNPs demonstrated negligible change in size under standard −20°C storage conditions. While empty MNPs were only stable at room temperature for one day, and not at 37°C, dye-loaded nanoparticles were stable for at least eight days under both storage conditions. We then performed *in vitro* studies to evaluate MNP cellular uptake mechanisms using the human renal cell carcinoma cell line 786-O treated with pharmacological inhibitors of uptake pathways. We found that MNP internalization is almost entirely prevented by dynamin inhibitors, while macropinocytosis inhibition also reduced uptake, suggesting that such standard nanoparticle uptake pathways are robust to the mesoscale size range. These findings provide key insights into the stability profile and endocytosis mechanisms of MNPs, which are critical for materials scale-up and translation of novel kidney-targeted drug and gene therapies.

## 1. Introduction

Over the past decade, nanomedicine-based gene therapy formulations have seen growing clinical success [1, 2]. Notably, Alnylam Pharmaceuticals developed small interfering RNA (siRNA) lipid nanoparticle (LNP) therapies such as Patisiran, which received U.S. Food and Drug Administration (FDA) approval in 2018 [3]. More recently, messenger RNA (mRNA) LNPs gained worldwide attention as the basis for SARS-CoV-2 vaccines, granted emergency use authorization by the FDA in 2020. Beyond vaccines, mRNA therapeutics continue to show promising potential in both pre-clinical and clinical settings [4]. Currently, there are around 100 approved nanomedicines on the market, and more than 600 in clinical trials, addressing a variety of disease areas such as cancer, infectious diseases, immunological diseases, and neurodegeneration [5, 6].

Our prior studies developed anionic mesoscale nanoparticles (MNPs), which are 300 – 500 nm in diameter and composed of diblock copolymers of poly(lactic-*co*-glycolic acid) conjugated to polyethylene glycol (PLGA-PEG) [7, 8]. MNPs demonstrate remarkable kidney-targeting ability, with up to 26-fold higher accumulation in the kidneys than in other organs. Specifically, MNPs localize primarily to renal tubular epithelial cells through transcytosis across the peritubular endothelium [7]. More recently, we encapsulated small-molecule dyes and reactive oxygen species scavengers within these MNPs [7, 9]. We also extended this platform to encapsulate siRNA, mRNA, and peptides [10-15].

Despite the demonstrated functionality of MNPs, there is still much to understand regarding the physicochemical parameters and how they control biological interactions. Environmental factors such as product transportation and storage could contribute to nanoparticle degradation, therefore impacting the safety, efficacy, and quality of nanomedicines [16, 17]. Studying these factors aid in identifying storage conditions that minimize degradation and maintain MNP drug product quality throughout shelf-life [16]. As MNPs, with a diameter of 300-500 nm, are larger than standard nanoparticle formulations of less than 200 nm, it is also key to understand their cellular uptake mechanisms. While prior studies have shown their ability to reach and accumulate in proximal tubular epithelial cells *in vivo* [7, 8], the specific endocytic pathways involved have not yet been determined. Understanding these cellular uptake pathways of MNPs is fundamental to advancing translation of kidney-targeting therapeutics, as the route of entry influences intracellular trafficking, release of the therapeutic payload, and ultimately, biological efficacy.

In general, nanoparticle uptake into cells is widely studied, including through such mechanisms as phagocytosis, clathrin-mediated, caveolin-dependent, clathrin/caveolae independent, macropinocytosis, and others [18]. Physicochemical properties of nanoparticles, including size, shape, surface charge, hydrophobicity, and surface functionality, can affect their cellular uptake by influencing the physical interaction between the nanoparticles and cell membrane [18, 19]. For instance, cell membrane permeability and integrity depends largely on the size and surface chemistry of interacting nanoparticles [20]. Size-dependent nanoparticle internalization studies revealed that nanoparticles with a size range of 120-150 nm are internalized via clathrin- or caveolin-mediated endocytosis pathways [21, 22]. In the caveolae-mediated pathway, the size of caveolae hinders the uptake of larger nanoparticles [23, 24]. In contrast, nanoparticle uptake mechanisms shift from clathrin- to caveolae-mediated with increasing size, which became the predominant entry pathway of nanoparticles with a size of 500 nm [22]. In addition, macropinocytosis has been shown to internalize nanoparticles ranging from a few hundred nanometers to a micrometer in size [25-27]. Different studies aimed at investigating the relationship between nanoparticle size and uptake mechanisms are still inconsistent, in part due to variance in materials [22, 25, 26]. PEGylated lipid nanomaterials demonstrated strong preference for clathrin- and caveolae-mediated endocytosis in human embryonic kidney cells [28]. Furthermore, 100 and 130 nm particles with high density PEG coating internalized via clathrin-mediated endocytosis, whereas 100 nm particles with low density PEG coating internalized via macropinocytosis [27]. Despite the interest and important, no studies have yet investigated the uptake of high-density PEG-coated mesoscale nanoparticles.

In this work, we sought to understand stability of MNPs and how their physicochemical properties interact with cellular biology. Specifically, we investigated particle stability under temperature stress conditions, as well as the cellular mechanisms that drive MNP internalization. These studies are necessary to further the clinical development of kidney-targeted therapies, while they are unique in that few others have explored these parameters within this size range of particle formulation.

## 2. Methods

### 2.1. PLGA conjugation to PEG

MNPs were synthesized from poly(lactic-*co*-glycolic acid) (PLGA) and polyethylene glycol (PEG). Briefly, 5 g of PLGA-COOH (Resomer RG504H; Sigma Aldrich, St. Louis, MO, USA; Cat. No. 719900-5G) was dissolved in 10 mL of dichloromethane (Sigma Aldrich; Cat. No. 270997) and reacted with 135 mg of N-hydroxysuccinimide (Sigma Aldrich; Cat. No. 130572-100g) and 230 mg of 1-ethyl-3-(3-dimethylaminopropyl)carbodiimide (Sigma Aldrich, Cat. No. E6383-1G) under stirring overnight to form PLGA-NHS. The polymer was then precipitated with 10 mL of cold ethyl ether (Thermo Fisher Scientific, Waltham, MA, USA; Cat. No. E138-1), washed with a cold ethyl ether:methanol mixture, vacuum-dried, and stored at −20°C until further use. PLGA-NHS was then dissolved in 4 mL of chloroform (Thermo Fisher Scientific; Cat. No. C606-4) and reacted with 250 mg of NH_2_-PEG-COOH (Nanocs, New York, NY, USA; Cat. No. PG2-AMCA-5K) in the presence of 37.7 µL of N,N-diisopropylethylamine (Sigma Aldrich; Cat. No. D125806) under overnight stirring. The resulting PLGA-PEG conjugate was precipitated into cold methanol to remove excess PEG, washed repeatedly with cold methanol, vacuum-dried, and stored at −20°C.

### 2.2. MNP formulation

Standard MNP formulation procedures were followed similarly to our previously established nanoprecipitation protocol [12]. Empty MNPs were formulated by dissolving 100 mg of PLGA-PEG co-polymer in 2 mL of acetonitrile with a stir bar and mixed on a stir plate for 10 minutes. Separately, 4 mL of purified water containing 75 μL of Pluronic F-68 (MP Biomedicals, Irvine, CA, USA; Cat. No. 092750049) was prepared under continuous stirring in a round-bottom flask. The organic phase containing the co-polymer was then introduced dropwise into the aqueous phase using a syringe pump at a controlled rate of 0.1 mL/min to facilitate nanoprecipitation. The reaction mixture was stirred for an additional 2 hours to allow solvent evaporation and nanoparticle hardening. The resulting empty MNP suspension was centrifuged at 7356 x g for 15 minutes at 4°C, and the pellet was washed three times with purified water to remove residual surfactant. The washed nanoparticles were resuspended in 10 mL of 2% (w/v) sucrose (Sigma Aldrich, Cat. No. S0389-500g) solution as a cryoprotectant, frozen at −80°C for at least one hour, then lyophilized for 48 hours.

For DEDC-loaded MNPs, 100 mg of PLGA-PEG co-polymer and 10 mg of 3,3′-Diethylthiadicarbocyanine iodide (DEDC) dye (Acros; Cat. No. 407470010) were added to 2 mL of acetonitrile with a stir bar and mixed on a stir plate for 10 minutes. The subsequent aqueous phase preparation and nanoprecipitation procedure are identical to the empty MNP formulation.

IL-6 siRNA (Horizon Discovery, Waterbeach, UK; Cat. No. L-043739-00-0005) and siGLO (Horizon Discovery, Cat. No. D-001630-02-05) were resuspended per the vendor’s protocol to a final amount of 5 nmol at a 10 µM stock concentration. IL-6 siRNA- and siGLO-loaded MNPs were formulated by dissolving 100 mg of PLGA-PEG co-polymer in 2 mL of acetonitrile containing either 10 µL of 10 µM IL-6 siRNA (for a total amount of 1 nmol siRNA) or 10 µL of 10 µM siGLO (for a total amount of 1 nmol siGLO), followed by bath sonication for 2 minutes. Separately, 4 mL of purified water containing 75 μL of Pluronic F-68 (MP Biomedicals; Cat. No. 092750049) was prepared under continuous stirring in a round-bottom flask. The organic phase containing the co-polymer and siRNA/siGLO was then introduced dropwise into the aqueous phase using a syringe pump at a controlled rate of 0.1 mL/min to facilitate nanoprecipitation. The reaction mixture was stirred for an additional 2 hours to allow solvent evaporation and nanoparticle hardening. The resulting siRNA/siGLO-loaded MNP suspension was centrifuged at 7356 x g for 15 minutes at 4°C, and the pellet was washed three times with purified water to remove unencapsulated siRNA/siGLO and residual surfactant. The washed nanoparticles were resuspended in 10 mL of 2% (w/v) sucrose (Sigma Aldrich, Cat. No. S0389-500G) solution as a cryoprotectant, frozen at −80°C for at least one hour, then lyophilized for 48 hours.

At the end of the lyophilization process, all MNPs were stored at −20°C until further use.

### 2.3. MNP physicochemical characterization

MNP physicochemical characteristics were analyzed to determine their size distribution, surface charge, and siRNA encapsulation. The hydrodynamic diameter and polydispersity index (PDI) of the MNPs were measured using dynamic light scattering (DLS) on a NanoZS90 Zetasizer (Malvern Panalytical, Malvern, United Kingdom), while ζ-potential was determined through electrophoretic light scattering to assess surface charge. The siRNA encapsulation efficiency of the MNPs was quantified using the Quant-iT RiboGreen RNA assay (ThermoFisher Scientific, Cat. No. R11490). Briefly, approximately 5 mg of MNPs were dissolved in acetonitrile and treated with 1X Tris-EDTA buffer to release the encapsulated siRNA, followed by centrifugation at max speed to collect the supernatant which contained released siRNA. Manufacturer protocol was then followed to quantity siRNA. The encapsulation efficiency (%) was determined based on the ratio of total encapsulated siRNA to the input siRNA initially used in the MNP formulation.

### 2.4. Nanoparticle stability studies

To evaluate freeze-thaw stability of empty and DEDC MNPs, each were resuspended in 1X PBS at 10 mg/mL. They were then subjected to repeated freeze-thaw, up to 4 cycles, and measured for hydrodynamic diameter and PDI by DLS. Upon resuspension in 1X PBS, MNPs were immediately measured for hydrodynamic diameter and PDI, which corresponded to the freeze-thaw cycle 0 group. Resuspended MNPs were then stored at −20°C under frozen conditions for at least 18 hours. Afterwards, frozen MNPs were thawed at room temperature (RT) then measured for hydrodynamic diameter and PDI, which corresponded to the freeze-thaw cycle 1 group. Then, MNP samples were stored again at −20°C under frozen condition for at least 18 hours. This freeze-thaw cycle process is repeated for up to 4 times.

To evaluate the stability of empty and DEDC MNPs outside the intended storage condition of −20°C, 10 mg of lyophilized empty MNPs were stored at RT, 37°C, and −20°C for 0, 1, 3, 7, 15, and 35 days, and DEDC MNPs were stored at RT, 37°C, and −20°C for 0, 2, 4, 8, 16 days. At each timepoint, empty and DEDC MNPs were resuspended in 1X PBS at 10 mg/mL and measured for hydrodynamic diameter and PDI.

### 2.5. Inhibition of MNP endocytosis in kidney cells

To evaluate siRNA-MNP endocytosis, we used fluorescently-labeled siRNA mimic siGLO MNPs with renal cell adenocarcinoma 786-O cells (American Type Culture Collection, Cat. No. CRL-1932). To inhibit clathrin-mediated endocytosis, 786-O cells were treated with 30 µM PitStop2 (abcam; Cat. No. ab120687) for 30 mins (short treatment) followed by either a 3X wash with 1X PBS or a no wash step (long treatment). Cells were then incubated with siGLO MNPs for 3 hours. Similarly, to inhibit dynamin-mediated endocytosis, 786-O cells were treated with 80 µM Dynasore (MilliporeSigma; Cat. No. 324410-10MG) for 30 mins (short treatment) followed by either a 3X wash with 1X PBS or a no wash step (long treatment), followed by siGLO MNP incubation for 3 hours. To inhibit macropinocytosis, 786-O cells were treated with 10 µM amiloride (Thermo Fisher Scientific; Cat. No. A25995G) for either 1 hour (short treatment) or 3 hours (long treatment), followed by siGLO MNP incubation for 3 hours. At the end of the 3 hour siGLO MNP treatment, cells were washed 3-times with 1X PBS to remove unbound siGLO MNPs then fixed with 10% formalin and stained with Hoechst 33342 (Thermo Fisher Scientific, Cat. No. 62249) and CellMask™ (Thermo Fisher Scientific; Cat. No. C37608) for nuclear and plasma membrane staining, respectively. Fixed cells were then fluorescently imaged using an EVOS M5000 Microscope (Thermo Fisher Scientific; Cat. No. AMF5000SV) for siGLO and nuclear quantification.

Fluorescence microscopy images were analyzed using ImageJ (NIH, Bethesda, MD, USA) [29]. Hoechst-stained nuclei were used to identify and count individual cells, while fluorescence corresponding to the siGLO cargo was used to quantify intracellular nanoparticle uptake. Regions of interest were drawn around each field, and the mean fluorescence intensity was measured via 585/628 nm excitation/emission. The total fluorescence intensity per image was then normalized by the corresponding cell count, yielding the mean fluorescence intensity per cell (MFI/cell). All images were processed using identical exposure settings and analysis parameters to ensure consistency across samples.

### 2.6. Flow cytometry

Flow cytometry (Attune NxT; Thermo Fisher Scientific) was used to quantify cellular uptake of siGLO encapsulated within MNPs and determine uptake efficiency (%siGLO+ cells). Briefly, siGLO MNP treated cells were gently washed 3-times with 1X PBS to remove unbound MNPs. Cells were then trypsinized using TrypLE (Thermo Fisher Scientific, Cat. No. 12604021) and neutralized with equal volume of complete RPMI media. siGLO fluorescence was detected in the yellow YL2 channel (excitation/emission 561/620 nm). Gating was performed sequentially as follows: first, side scatter-height (SSC-H) versus forward scatter (FSC-H) was used to identify the main cell population and exclude debris; next, SSC-H versus SSC-A (side scatter-area) was applied to gate singlets and remove doublets; finally, fluorescence intensity in the YL2-H channel was plotted to distinguish siGLO+ cells. The negative control (untreated cells) was used to define the fluorescence threshold for siGLO+ cells. Data were analyzed using Attune NxT software to calculate %siGLO+ cells and the corresponding mean fluorescence intensity.

### 2.7. In vitro IL-6 expression and siRNA-MNP knockdown

786-O cells were seeded at 40,000 cells/well in a 96 well plate and incubated for 24 hours at 37°C, 5% CO_2_ using growth medium consisting of HyClone RPMI 1640 (Cytiva, Marlborough, MA, USA; Cat. No. SH30255FS) + 10% heat-inactivated fetal bovine serum (HI-FBS) (Thermo Fisher Scientific; Cat. No. MT35011CV) + 1X Primocin (Invivogen, San Diego, CA, USA; Cat. No. ANT-PM-1). Cells were then treated with lipopolysaccharide (LPS) from *Escherichia coli* O111:B4 (Millipore Sigma; Cat. No. L2630) at 20 µg/mL to induce Interleukin-6 (IL-6) expression. Induced cells were treated with 2.74, 13.7, and 27.4 ng of IL-6 siRNA encapsulated within MNPs. Cells were then incubated for 48h under standard conditions.

Total RNA was extracted using the PureLink RNA Mini Kit (Thermo Fisher Scientific, Cat. No. 12183018A) according to the manufacturer’s protocol. Cells were lysed in lysis buffer containing β-mercaptoethanol (10 µL for each 1 mL of lysis buffer), and the lysates were homogenized by pipetting. The clarified lysate was mixed with ethanol and loaded onto the spin cartridge, followed by centrifugation to allow RNA binding to the silica membrane. The column was then washed per manufacturer instructions to remove residual contaminants. RNA was eluted in RNase-free water after a final spin. On-column DNase treatment (PureLink™ DNase Set, Thermo Fisher Scientific) was included to remove genomic DNA. RNA quantity and purity were determined using the A260/A280 ratio.

Complementary DNA (cDNA) was synthesized from total RNA using the High-Capacity cDNA Reverse Transcription Kit (Applied Biosystems, 4375575) according to the manufacturer’s protocol. Briefly, 2 µg of purified RNA from *in vitro* lysates was combined with a 2X reverse transcription master mix containing random primers, dNTPs, MultiScribe reverse transcriptase, and RNase inhibitor. Reactions were gently mixed, briefly spun down, and incubated in a thermal cycler at 25°C for 10 min, 37°C for 120 min, and 85°C for 5 min, followed by holding at 4°C. The resulting cDNA was stored at −20°C until use for downstream RNA expression analyses.

qPCR was performed using the TaqMan Fast Advanced Master Mix (Applied Biosystems; Cat. No. 4444557) following the manufacturer’s instructions (TaqMan Master Mix User Guide, Thermo Fisher Scientific). Prior to use, all reagents were mixed thoroughly and TaqMan Gene Expression Assays were thawed on ice and protected from light. Each 20 µL reaction contained 10 µL of TaqMan Master Mix, 1 µL of human IL-6 FAM-labeled target assay (Thermo Fisher Scientific; Cat. No. Hs00174131_m1), 1 µL of human GAPDH VIC-labeled endogenous control assay (Thermo Fisher Scientific; Cat. No. Hs02786624_g1), 2 µL of cDNA template, and 6 µL of nuclease-free water. No-template controls were included for each assay. Reactions were run in triplicate on an Applied Biosystems real-time PCR system under standard cycling conditions (95°C for 20 s, followed by 40 cycles of 95°C for 1 s and 60°C for 20 s). Relative gene expression levels were calculated using the ΔΔCt method, normalizing IL-6 expression to GAPDH housekeeping gene.

Secreted human IL-6 protein levels in cell culture supernatants were measured using the Invitrogen Human IL-6 ELISA Kit (Thermo Fisher Scientific, Cat. No. EH2IL6) according to the manufacturer’s instructions. Briefly, 50 µL of standards or samples were added to pre-coated wells followed by 50 µL of biotinylated anti IL-6 antibody reagent. After a 2-hour incubation at room temperature, wells were washed three times and incubated with 100 µL of streptavidin-HRP solution for 30 minutes. Following three additional washes, 100 µL of TMB substrate was added and the reaction was stopped after 30 minutes with 100 µL of stop solution. Absorbance was read at 450 nm using a microplate reader (BioTek Synergy; Agilent Technologies, Santa Clara, CA, USA). IL-6 concentrations were calculated from the standard curve generated in parallel.

### 2.8. Statistical analysis

All statistical analyses were conducted in R (version 4.3.0) using the packages car, emmeans, multcomp, sandwich, and ggplot2. Data were first examined for normality (Shapiro-Wilk test) and homogeneity of variances (Levene’s test). One-way analysis of variance (ANOVA) was performed with *post hoc* Dunnett’s test to compare each treatment group against the corresponding control. For experiments with multiple treatment doses or timepoints, one-way ANOVA followed by Dunnett-adjusted p-values was applied. For interaction analyses, a two-way ANOVA was performed using linear models, with pairwise contrasts estimated via estimated marginal means (EMMs) and heteroskedasticity-consistent (HC3) robust covariance to ensure reliability under variance heterogeneity. Results were reported as mean ± SEM, and differences were considered statistically significant at *p* < 0.05. Graphs were generated using ggplot2 in R or GraphPad Prism.

## 3. Results and Discussion

### 3.1. MNP formulation

To perform stability, uptake, and functional knockdown studies, we separately formulated MNPs that encapsulate cargos 3,3′-Diethylthiadicarbocyanine iodide (DEDC), IL-6 siRNA, and siGLO (a fluorescent oligonucleotide duplex), as well as empty MNPs as control. Empty MNPs had an average hydrodynamic diameter of 287.0 ± 3.5 nm, a polydispersity index (PDI) of 0.26 ± 0.03, a ζ-potential of −19.2 ± 0.4 mV. DEDC MNPs were 306.3 ± 9.6 nm, with a PDI of 0.18 ± 0.04 and a ζ-potential of −24.2 ± 1.2 mV. IL-6 siRNA MNPs demonstrated an average hydrodynamic diameter of 357.9 ± 13.1 nm, a PDI of 0.22 ± 0.04, a ζ potential of −20.9 ± 0.3 mV, and an encapsulation efficiency of 53.3%. siGLO MNPs were 314.7 ± 3.1 nm, with a PDI of 0.11 ± 0.04 and a ζ-potential of −32.9 ± 0.5 mV, and an encapsulation efficiency of 47.4%. Overall, these results indicate the versatility of the MNP formulation platform in achieving consistent anionic nanoparticle formulations in the mesoscale range.

**Table 1.** Physicochemical characteristics of MNPs with cargos encapsulated as indicated. Data reported are average of 3 technical measurements. SD: standard deviation; PDI: polydispersity index; ζ potential; EE: encapsulation efficiency.

| | Size ± SD (nm) | PDI ± SD | $\zeta \pm$ SD (mV) | EE (%) | Loading (ng/mg particle) |
| --- | --- | --- | --- | --- | --- |
| **Empty MNPs** | $287.0 \pm 3.5$ | $0.26 \pm 0.03$ | $-19.2 \pm 0.4$ | N/A | N/A |
| **DEDC MNPs** | $306.3 \pm 9.6$ | $0.18 \pm 0.04$ | $-24.2 \pm 1.2$ | N/A | N/A |
| **IL-6 siRNA MNPs** | $357.9 \pm 13.1$ | $0.22 \pm 0.04$ | $-20.9 \pm 0.3$ | 53.3 | 27.4 |
| **siGLO MNPs** | $314.7 \pm 3.1$ | $0.11 \pm 0.04$ | $-32.9 \pm 0.5$ | 47.4 | 26.2 |

### 3.2. MNPs are stable for up to eight days at room temperature and can withstand multiple freeze-thaw cycles

We found that exposing empty MNPs for up to 4 freeze-thaw cycles had a negligible effect on the hydrodynamic diameter (ranging between 283.8 – 293.9 nm) and PDI (ranging between 0.17 – 0.19) when compared to empty MNPs prior to freeze-thaw cycles (freeze-thaw cycle 0), with a mean hydrodynamic diameter of 283.8 nm and PDI of 0.16) (Figure 1A). Similarly, exposing DEDC MNPs for up to 3 freeze-thaw cycles also had a negligible effect on the hydrodynamic diameter (ranging between 313.3 – 332.4 nm) and PDI (ranging between 0.16 – 0.22) when compared to DEDC MNPs prior to freeze-thaw cycles, with a mean hydrodynamic diameter of 326.0 nm and PDI of 0.21 (Figure 1B). These results indicate that MNPs can withstand multiple freeze-thaw cycles without significant aggregation. Further *in vitro* and *in vivo* efficacy studies are required on these freeze-thawed MNPs to ensure they retain their efficacy.

**Figure 1.**
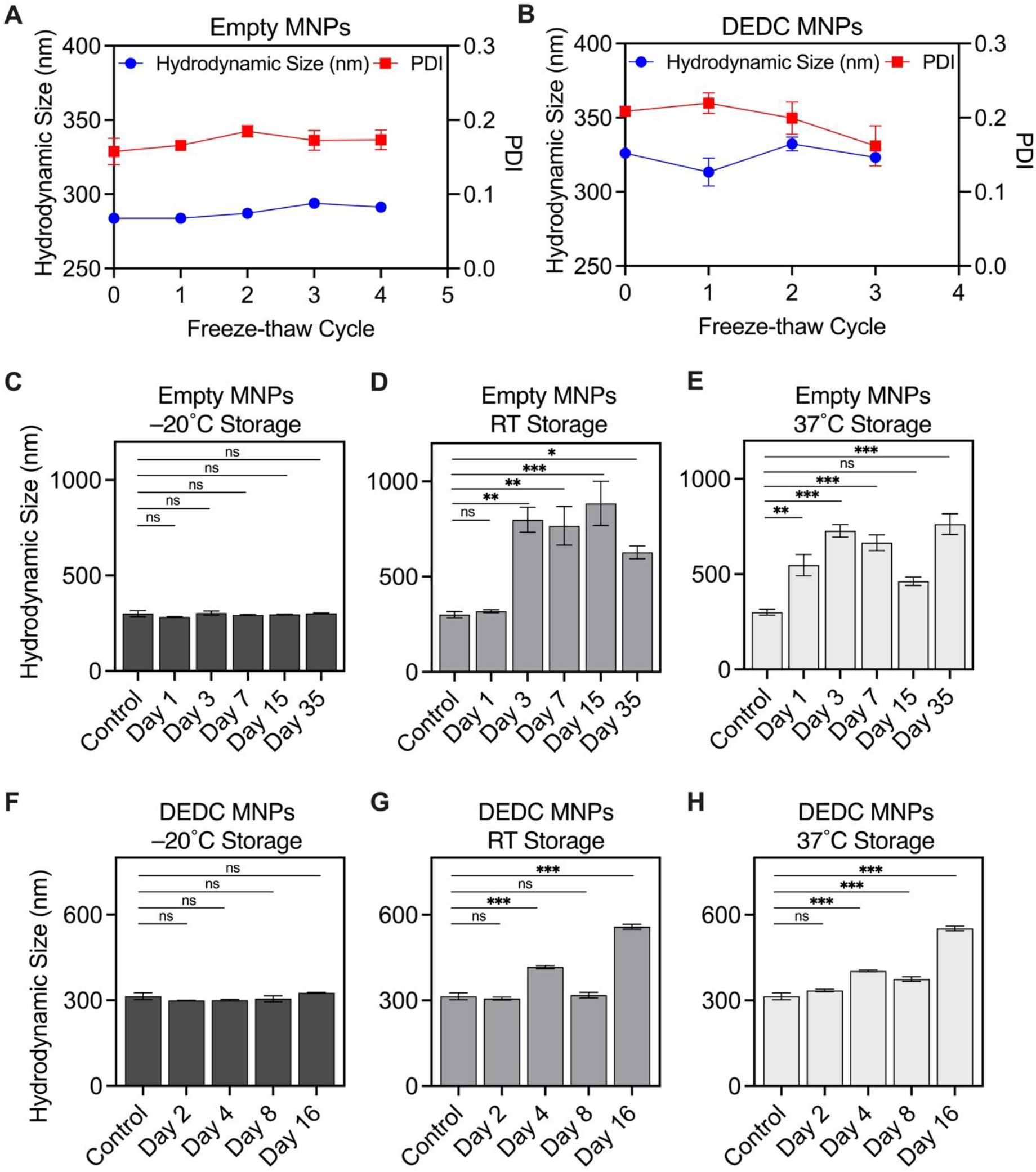
MNPs are stable at −20°C and can withstand multiple freeze-thaw cycles. A) Hydrodynamic diameter and PDI of empty MNPs under successive freeze-thaw cycles. B) Hydrodynamic diameter and PDI of DEDC MNPs under successive freeze-thaw cycles. C) Hydrodynamic diameter of empty MNPs held at −20°C for up to day 35. D) Hydrodynamic diameter of empty MNPs held at RT for up to day 35. E) Hydrodynamic diameter of empty MNPs held at 37°C for up to day 35. F) Hydrodynamic diameter of DEDC MNPs held at −20°C for up to day 16. G) Hydrodynamic diameter of DEDC MNPs held at RT for up to day 16. H) Hydrodynamic diameter of DEDC MNPs held at 37°C for up to day 16. RT: room temperature. Data shown are mean ± SD of 3 independent replicates.

Time out of intended storage was evaluated by exposing lyophilized empty and DEDC MNPs to room temperature (RT; 22 – 24°C) and 37°C for prolonged periods of time, in addition to the −20°C control group. Empty MNPs showed no statistically significant change in hydrodynamic diameter when stored at −20°C for up to 35 days, with mean MNP hydrodynamic diameter ranging between 284 – 302.4 nm (Figure 1C). When incubated at RT, empty MNPs demonstrated an increased hydrodynamic diameter of 798.8 nm at 3 days (*p* < 0.01), 766.8 nm at 7 days (*p* < 0.01), 884.5 nm at 15 days (*p* < 0.001), and 627.3 nm at 35 days (*p* < 0.05) compared to control (300.5 nm), whereas the 1-day group did not differ significantly (318.3 nm) when compared to the control (Figure 1D). Similarly, when incubated at 37°C, empty MNPs demonstrated an increased hydrodynamic diameter of 547.6 nm at 1 day (*p* < 0.01), 727.6 nm at 3 days (*p* < 0.001), 665.3 nm at 7 days (*p* < 0.001), and 762.5 nm at 35 days (*p* < 0.001) compared to control, whereas the 15-day (462.6 nm) group did not differ significantly (Figure 1E). DEDC MNPs showed no change in hydrodynamic diameter when stored at −20°C for up to 15 days, with mean diameter ranging between 300.5 – 326.0 nm. When incubated at RT, DEDC MNP size increased significantly to 416.7 nm at 4 days (*p* < 0.001), though still within the mesoscale, and 558.4 nm at 16 days (*p* < 0.001), whereas the 2-day (306.3 nm) and 8-day (318.2 nm) groups did not differ significantly to the control (314.2 nm) (Figure 1G). When incubated at 37°C, DEDC MNP diameter increased significantly compared to controls with a mean diameter of 403.2 nm at 4 days (*p* < 0.001) 375 nm at 8 days (*p* < 0.001), both within the mesoscale, and 552.4 nm at 16 days (*p* < 0.001), whereas the 2-day group did not differ significantly, with a mean diameter of 334.7 nm (Figure 1H). However, in all cases prior to 16 days, increases were minimal, and still within the mesoscale range, which was not true of empty MNPs. These results indicate that cargo appears to stabilize MNPs, and when done with this fluorescent dye, they are stable for at least 8 days out of the freezer. This is supported by other studies that demonstrate particle stability is improved by cargo loading[30]. However, we can hypothesize that the presence of encapsulated cargo could alter the internal organization of the PLGA core, reducing polymer chain mobility and limiting time-dependent rearrangements that may otherwise promote aggregation in empty MNPs. These findings underscore the importance of characterizing payload distribution in nanoparticle formulations, as a higher proportion of empty nanoparticles may compromise the stability of the final drug product and reduce its therapeutic effect [31].

### 3.3. siGLO MNPs show efficacy in 786-O cells

786-O cells were stained with Hoechst 33342 nuclear stain and CellMask plasma membrane stain, then incubated with siGLO MNPs for 2 hrs. Microscopy images reveal a combination of punctate and diffuse siGLO fluorescence signal that are localized intracellularly (Figure 2A). We also used flow cytometry to measure uptake, with gating to exclude debris and doublets, and no-fluorescence controls used to set a boundary for positive uptake (Figure 2B). 786-O cells were treated with 0, 10, 20, 40, and 80 ng of siGLO encapsulated in MNPs (also equivalent to 0, 3.6, 7.3, 14.6, and 29.2 mg/mL MNPs), demonstrating a dose-dependent increase in signal intensity. On average, 24.8% of cells treated with 10 ng of siGLO within MNPs exhibited fluorescence above the threshold, 44.8% of cells for 20 ng siGLO treatment, 65.9% of cells for 40 ng siGLO treatment, and 76.2% of cells for 80 ng siGLO treatment (Figure 2C and 2D).

**Figure 2.**
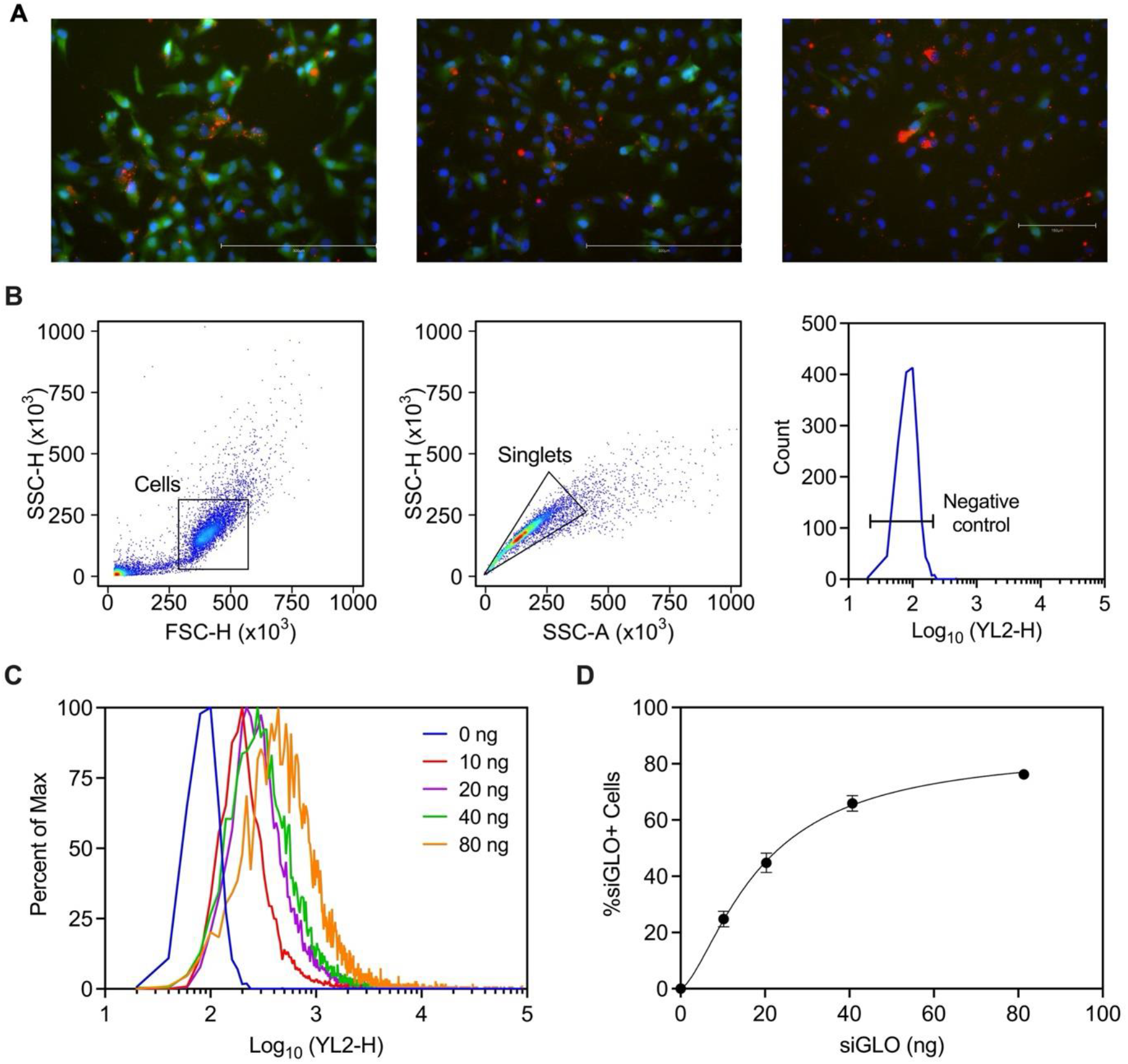
siGLO MNPs are taken up by 786-O cells. A) Microscopy images of 786-O cells upon treatment with siGLO MNPs. Red: siGLO; Green: Cell Membrane; Blue: Nucleus. B) Flow cytometry gating strategy. C) Fluorescence signal distribution after treatment with increasing siGLO concentrations encapsulated within MNPs. D) Percent of cells with fluorescence above 0 ng siGLO controls. Scale bar: 300 µm.

These results demonstrate successful cellular uptake of siGLO MNPs, as evidenced by the cytoplasmic expression of siGLO in red (Figure 2A). These results corroborate our previous study that showed successful uptake of mCherry mRNA MNPs in 786-O cells [12]. Interestingly, fluorescence microscopy images also revealed both a punctate and diffuse siGLO fluorescence, suggesting a combination of MNP entrapment within cellular compartments and subsequent release into the cytoplasm (Figure 2A). Endosomal escape of nanoparticles is a well-known bottleneck in nanoparticle-mediated delivery [32, 33]. Previous electron microscopy studies using gold particle-labeled siRNA have shown that only approximately 2% of particles are released into the cytoplasm, while the majority remain trapped within endosomal vesicles [34]. The endosomal escape of MNPs hasn’t been studied yet. Future work focused on understanding the endosomal escape mechanisms and rate of MNPs will be essential for understanding their clinical utility. Finally, microscopy-based observations of cellular uptake were corroborated by flow cytometry, revealing a dose-dependent uptake (Figure 2C and 2D).

### 3.4. MNPs are predominantly internalized via dynamin-dependent endocytosis with secondary contributions from macropinocytosis

As we determined that MNPs are taken up by 786-O cells, we sought to elucidate which endocytosis pathways are involved in MNP cellular uptake. We investigated three known uptake pathways, clathrin-mediated, dynamin-mediated, and macropinocytosis-mediated endocytosis by using pharmacological inhibitors for each endocytosis pathway.

We used Pitstop2 to inhibit clathrin-mediated endocytosis, as it interferes with binding to its N-terminal domain [35]. We found no statistically significant difference between the PitStop2 short treatment group (30 mins inhibitor + Wash) and the control group with no inhibitor at all the siGLO doses tested (Figure 3A). However, in the PitStop2 long treatment (30 mins pre-incubation with inhibitor, 2 hr with MNPs + inhibitor), we observed a 3.2-fold decrease (*p* < 0.001), 1.5-fold decrease (*p* < 0.0001), and a 2.0-fold decrease (*p* < 0.0001) at the 2.6 ng, 13.1 ng, and 26.2 ng siGLO doses respectively (Figure 3A). These results indicate that clathrin inhibition via PitStop2 is reversible after inhibitor is washed out. This observation aligns with previous studies showing that clathrin-mediated endocytosis is fully restored within 30 minutes of inhibitor removal (short treatment) [36]. When cells were treated for a longer period, without washing out the inhibitor, we observed up to 3.2-fold inhibition of clathrin-mediated endocytosis (Figure 3A), indicating partial uptake of siGLO MNPs through the clathrin pathway.

**Figure 3.**
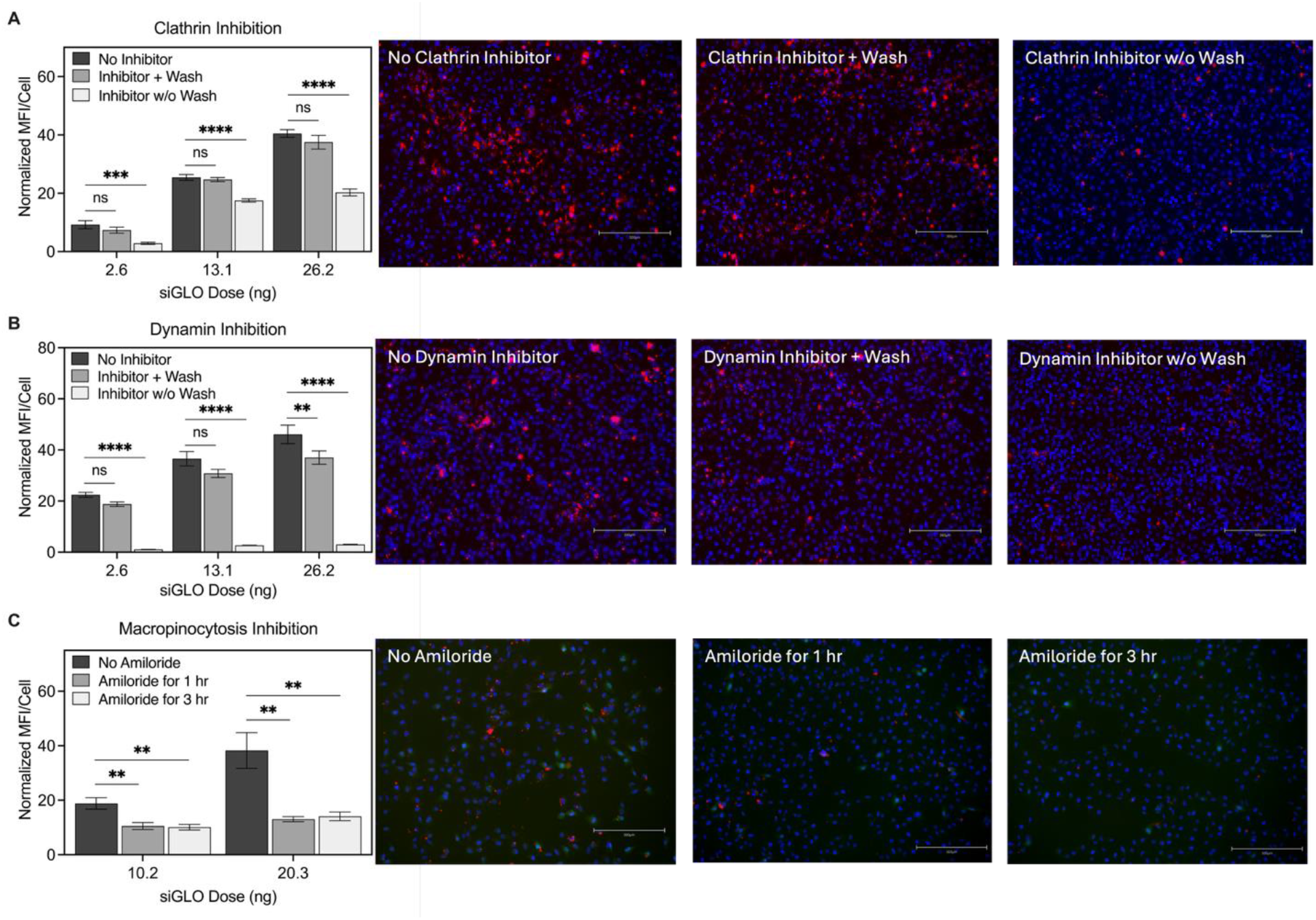
MNPs are predominantly internalized via dynamin-dependent endocytosis with secondary contributions from macropinocytosis. A) clathrin inhibition using 30 µM PitStop2 for 30 mins followed by wash or no wash during MNP incubation. B) dynamin inhibition using 80 µM Dynasore for 30 mins with wash or for 30 mins without washing. C) macropinocytosis inhibition using 10 µM Amiloride for either 1 hour or 3 hours. No inhibitor was included as a control group. Scale bar: 300 µm.

To inhibit dynamin-mediated endocytosis, we used Dynasore, which inhibits dynamin by blocking its GTPase activity [37]. There was no statistically significant difference between the Dynasore short treatment group and the no-inhibitor control group at 2.6 ng and 13.1 ng siGLO doses tested (Figure 3B). At the 13.1 ng siGLO dose, there was a statistically significant difference between the Dynasore short treatment group and the no inhibitor group (*p* < 0.01), as shown by a minimal 1.2-fold decrease in siGLO expression (Figure 3B). In the Dynasore long treatment, we observed substantial decreases of 21.1-fold (*p* < 0.0001), 13.6-fold (*p* < 0.0001), and 15.3-fold (*p* < 0.0001) at the 2.6 ng, 13.1 ng, and 26.2 ng siGLO doses, respectively (Figure 3B). Similar to clathrin inhibition, dynamin inhibition via Dynasore is also reversible upon wash out as reported in previous studies [38]. However, when cells are treated for a longer period without washing out Dynasore and in the presence of siGLO MNPs, we observed a near complete dynamin-mediated endocytosis inhibition. Dynamin-dependent pathways include clathrin-, caveolin-, fast-endophilin-, RhoA-, and Arf6-associated endocytosis [39]. While our data reveal partial uptake through clathrin, further exploration of the additional dynamin-dependent routes would help delineate the specific contributions of each pathway to MNP internalization.

To inhibit macropinocytosis, we treated cells with amiloride, which inhibits Na^+^ channels and Na^+^/H^+^ exchange [40]. Macropinocytosis inhibition resulted in a 1.8-fold (*p* < 0.01) and 1.9-fold (*p* < 0.01) reduction in MNP uptake after 1 hour of amiloride incubation for 10.2 ng and 20.3 ng of siGLOP MNPs, respectively (Figure 3C). Similarly, a 2.9-fold (*p* < 0.01) and 2.7-fold (*p* < 0.01) reduction in MNP uptake was observed after 3 hours of amiloride incubation for 10.2 ng and 20.3 ng of siGLO MNPs, respectively (Figure 3C). These findings suggest that macropinocytosis contributes partially to MNP internalization. Prior work has indeed demonstrated that some particles larger than 200 nm may be internalized by macropinocytosis [39, 41].

Taken together, the endocytosis data indicate that MNPs are primarily internalized through dynamin-dependent pathways, with secondary involvement of macropinocytosis, a pathway known for internalization of larger particles. Pharmacological inhibitors remain valuable tools for studying nanoparticle internalization, but their use have several limitations, such as off-target effects [37, 42, 43] and a strong dependence on cell type [44]. To address these limitations, future studies could incorporate complementary approaches such as genetic knockdowns to help further confirm the role of specific endocytosis pathways.

### 3.5. MNPs encapsulating IL-6 siRNA knocked down IL-6 RNA and protein expression in 786-O cells

To establish a model of renal tubular inflammation, 786-O cells were treated with 0 (no LPS control), 1, 10, and 50 µg/mL of LPS and incubated for 48 hours to induce IL-6 mRNA expression (Figure 4A). Compared to the no LPS control, treatment with 1 µg/mL of LPS resulted in a significant 17.2-fold increase in IL-6 mRNA expression (*p* < 0.0001), 10 µg/mL of LPS led to a 14.7-fold increase (*p* < 0.0001), and 50 µg/mL of LPS produced a 23.9-fold increase (*p* < 0.0001) (Figure 4B). Cells were then treated with 0 (no MNPs + 20 µg/mL LPS), 2.74, 13.7, and 27.4 ng of IL-6 siRNA encapsulated within MNPs in the presence of 20 µg/mL of LPS and incubated for 48 hours. Relative to the LPS-only group, the 2.74 ng siRNA MNP treatment showed a slight but not significant decrease in IL-6 mRNA expression (*p* = 0.078), whereas the 13.7 ng and 27.4 ng siRNA MNP treatments resulted in significant 1.7-fold (*p* < 0.01) and 1.3-fold (*p* < 0.05) reductions, respectively (Figure 4C). Finally, compared to the LPS-only group, the 2.74 ng siRNA MNP treatment led to a significant 2.7-fold reduction in IL-6 protein expression (*p* < 0.05), the 13.7 ng treatment showed a 2.7-fold reduction (*p* < 0.05), and the 27.4 ng treatment showed a 2.4-fold reduction (*p* < 0.05) (Figure 4D).

**Figure 4.**
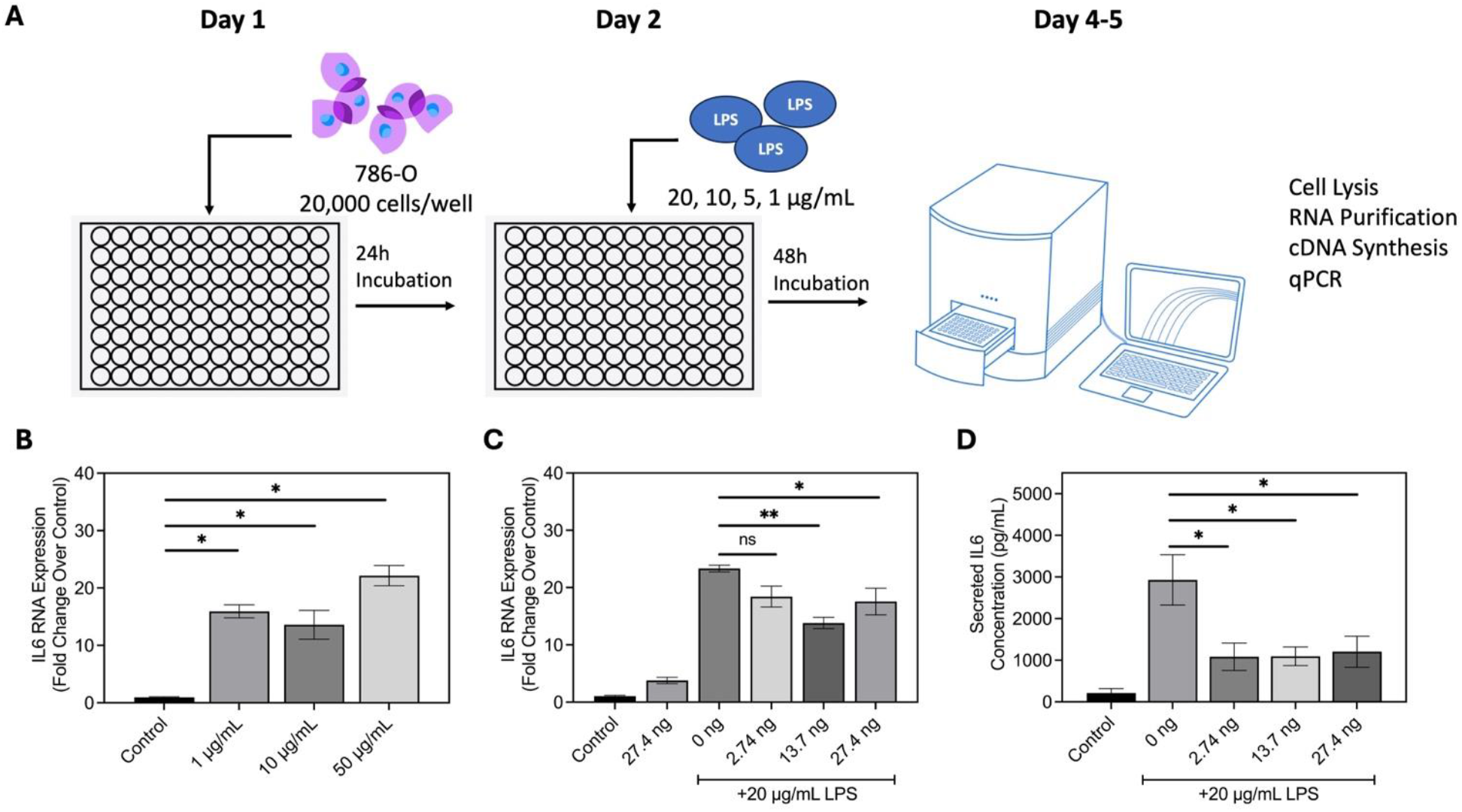
MNPs encapsulating IL-6 siRNA knocked down IL-6 mRNA and protein expression in 786-O cells. A) Illustration of the assay workflow for inducing IL-6 mRNA expression using LPS. B) IL-6 mRNA expression under various LPS concentrations; control: no LPS. C) IL-6 mRNA expression as a function of siRNA MNP dose and in the presence of 20 µg/mL LPS. D) Secreted IL-6 protein expression as a function of siRNA MNP dose and in the presence of 20 µg/mL LPS.

LPS is commonly used to induce acute kidney injury in animal models, primarily due to the septic inflammatory response it causes[45]. It is a key component of the outer membrane of Gram-negative bacteria and is widely used to stimulate cytokine production *in vitro* through activation of innate immune signaling pathways. LPS binds to toll-like receptor 4 (TLR4) on the cell surface, triggering downstream activation of NF-κB and IRF3 signaling cascades that drive transcription of pro-inflammatory cytokines, including IL-6 and IL-1β [46, 47]. We used LPS-induced IL-6 expression as a model to evaluate MNP-mediated siRNA knockdown efficacy. The mRNA knockdown we observed corresponded with a significant decrease in IL-6 protein expression. Interestingly, the dose-dependent knockdown observed at the mRNA level was not reflected in protein expression, where all doses produced comparable reductions. The reason for this lack of correlation is unclear but could be attributed to several factors. It is possible that even a modest reduction in IL-6 RNA is sufficient to substantially decrease protein production, beyond which additional RNA knockdown provides no further efficacy benefit. Alternatively, the slower turnover and translation rate of IL-6 protein compared to mRNA may contribute to this lack of correlation [48]. Time-course studies with extended incubation periods may help clarify the kinetics of IL-6 mRNA and protein regulation and provide deeper insight into this phenomenon.

In conclusion, we formulated MNPs that encapsulate both a small molecule dye as well as nucleic acids, such as IL-6 siRNA for therapeutic applications and siGLO for studying cellular uptake and trafficking. We also showed that dye-loaded MNPs remain stable at −20°C for long-term storage and both body and room temperature for up to 8 days, while they can withstand up to 4 freeze-thaw cycles. Additionally, MNPs are predominantly internalized through dynamin-mediated endocytosis with secondary contributions from macropinocytosis in 786-O cells. For future work, we plan to investigate additional dynamin-mediated pathways, such as caveolin- and fast endophilin-mediated endocytosis, as well as dynamin-independent pathways to further elucidate the contribution of these mechanisms to MNP internalization. Finally, examining these endocytosis pathways in primary tubular epithelial cells or endothelial cells will help uncover potential cell type-dependent differences in endocytosis.

## Acknowledgements

This work was supported by the NYCenter for Advancing Translational and Research Health (U54MD017979), Dialysis Clinic, Inc. Research Reserve Fund (#4231), and the SUNY Empire Innovation Program (Award #250010).

## Disclosures

AA is an employee of Eli Lilly and Company. RMW is the Founder of Zipcode Therapeutics, Inc.

## Notes

### Summary of Updates

The author list has been updated to include P. Ghosh.

